# Neural responses in the pain matrix when observing pain of others are unaffected by testosterone administration

**DOI:** 10.1101/245001

**Authors:** Sarah J. Heany, David Terburg, Dan J. Stein, Jack van Honk, Peter A. Bos

**Author notes:** Corresponding author: Peter A. Bos, PhD, Experimental Psychology, Utrecht University, Heidelberglaan 2, 3584 CS, Utrecht.

## Abstract

There is evidence of testosterone having deteriorating effects on cognitive and affective empathy. However, whether testosterone influences core affective empathy, that is empathy for pain, has not yet been investigated. Therefore, we tested neural responses to witnessing others in pain in a within-subject placebo controlled testosterone administration study. Using functional magnetic resonance imaging we provide affirming evidence that the empathy inducing paradigm causes changes in the activity throughout the pain circuitry, including the bilateral insula and anterior cingulate cortex. Administration of testosterone however did not influence these activation patterns in the pain matrix. Testosterone has thus downregulating effects on aspects of empathic behaviour, but based on these data does not seem to influence neural responses during core empathy for pain. This finding gives more insight into the role of testosterone in human empathy.

## 1 Introduction

The steroid hormone testosterone (T) is involved in a broad repertoire of human social-emotional behaviour(1), and plays a critical role in the sexual differentiation of such behaviour (2). For example, T administration studies have shown T to reduce fear responses (3), reduce trust in others (4), reduce submissive gazing (5), and increase aggressive responses to others (6). Overall, T seems to increase status seeking and dominance motivation (7), and facilitate behavioural responses dealing with threat or challenge (1). As such, several studies have shown T to reduce empathic behaviour. Hermans et al. (8) showed reduced facial imitation of the emotional expressions of others, which might reflect reduced emotion sharing, an aspect of affective empathy. T also reduces mindreading, the cognitive empathic ability whereby one infers the emotional state of the other from limited information. In women, two studies showed reduced performance on the Reading the Mind in the Eyes test after T (RMET,9,10), and a recent functional magnetic resonance imaging (fMRI) study shows this effect can depend on reduced neural connectivity of the frontal brain regions with the anterior cingulate cortex (ACC) and premotor regions (11). In men, T has also been shown to reduce RMET performance, although with the effect being dependent on a marker for fetal T exposure (the 2D:4D ratio) and psychopathic traits (12). Thus, whereas these studies show that T reduces the cognitive empathic ability of mindreading, as well the motor imitation of facial expressions, a core aspect of affective sharing -empathy for pain-has not yet been studied. Empathy for pain is an evolutionarily conserved emotional response that has been recorded in rodents (13). Indeed, responses of emotional state sharing have been noted consistently in mammals, and are thought to be crucial to survival of individuals (14).

A meta-analysis on the topic identified a core empathy for pain network comprising the bilateral insula and medial/anterior cingulate cortex, with other regions being additionally recruited depending on the functional paradigm being utilised, for example somatosensory regions were activated when the task used was picture based, and specifically, viewing body parts in pain recruited the inferior parietal/ventral premotor cortices (15). In another study the bilateral insula and rostral ACC were activated in response to receiving a painful stimulus, and also in response to the knowledge of a loved one being in pain, while the sensorimotor regions and caudal ACC were activated only when receiving pain directly (16). The sensorimotor cortex and SMA however were noted to have increased activity when viewing others in pain by Decety et al. (17) as well as the ACC and insula.

Although activation in the pain circuitry in response to seeing others in pain has repeatedly been observed and is considered a robust phenomenon, it has also been shown sensitive to contextual and endocrine manipulations. Several studies have found physiological responses to pain of others to be modulated by in/outgroup bias. fMRI studies have found decreased activation of the ACC (18) and bilateral anterior insula (19) when viewing outgroup members’ pain, and a TMS study noted a decreased empathic reaction in the corticospinal system when viewing outgroup pain (20). Hormonal factors can also affect empathy for pain, as the neuropeptide oxytocin has shown to decrease neural activation of the pain circuitry upon seeing pain in others, an effect that was independent of group status of the observed other (21).

The question to what extent T could alter neural responses to pain in others, and whether this would depend on ethnicity of the outgroup is currently unknown, and therefore in the current study we investigate the effect of T on neural empathic responses to pain in others. Based on the effects of T on cognitive empathy and facial mimicry, T might reduce such empathic responses. However, empathy for pain is different from mindreading or mimicry, as illustrated by contrasting findings from oxytocin administration studies showing increased cognitive empathy (e.g. 22), but reduced neural responses to pain in others (21). Effects of T might thus also show divergent effects on these different aspects of empathy.

## 2 METHODS

### 2.1 Participants

Thirty healthy young females were recruited to participate in the experiment. Ethical approval for human participant recruitment was granted by the Human Research Ethics Committee of the University of Cape Town, and the study was performed in accordance with the latest declaration of Helsinki. All participants provided written informed consent, and were screened for history of psychiatric conditions. Additional exclusion criteria were: current or recent use of psychotropic medications, use of hormonal contraceptives, pregnancy, abnormal menstrual cycle, any endocrine disorders, any other serious medical condition, left-handedness, habitual smoking, hearing problems, and colour blindness. After recruitment data of five participants were lost due to technical problems at the scanning facility, and one participant did not comply with the task instructions, leaving a final sample of 24 participants (age range 18-37, mean age = 21.2, SD = 4.19). Thirteen of the participants were black, three were of mixed ethnicity, and eight were white. In line with T administration literature (1), only women were considered as participants because the parameters, including quantity and time course, for inducing neurophysiological effects after a single sublingual administration of 0.5mg of T are known in women, whereas these parameters are not known in men (23). Each participant was paid ZAR250 (approx. €20) for their participation in this plus two other tasks unrelated to the current task.

### 2.2 Drug administration

Based on the existing literature on T administration, we closely followed the procedure reported by Tuiten et al (23). The procedure involves the sublingual administration of 0.5 mg of T with a hydroxypropyl-β-cyclodextrin carrier (manufactured by Laboswiss AG, Davos, Switzerland) to healthy young females. This is a well-established single dose T administration procedure that has been widely used and has been reliably shown to generate behavioural effects in young women (for a review, see (1)). For the dosage chosen, no side effects have been reported in this or other studies to date.

### 2.3 Procedure

Participants were tested on two separate days. The first and second sessions were separated by at least 48 hours. Both testing days fell within the follicular phase of the participants’ menstrual cycles, in order to ensure low and stable basal levels of sex hormones (e.g. progesterone, luteinizing hormone, and follicle stimulating hormone). On both occasions they arrived at the lab to receive the drug or placebo administration four hours before their scheduled MRI scan. Participants were instructed not to participate in any activity that may cause excessive fluctuations in hormone production, such as sports games, exams, or sexual activity before returning to the lab to undergo a MRI scan. This experiment was part of a larger study and all participants performed two other tasks during the scan session which were unrelated to the current task.

### 2.4 Empathy for pain task

In order to generate pain and no-pain conditions, short video clips of hands were shown to the participant. One hand at a time was viewed on screen in clips of 2.5 seconds duration. The viewed hand was either being injected by a hypodermic needle (test stimulus), or being gently prodded with an ear bud (control stimulus), as shown in Figure 1A. The hypodermic needle was fake and no hands were genuinely injected in the making of the task. The tip of the needle or ear bud made contact with the hand 1 second into each video clip. In between each video clip the participant saw a black fixation cross on a white background for between 3 and 8 seconds. Hands of two different colour were used; black hands and white hands. This resulted in four stimuli conditions (2 X 2; pain X colour) during the task. The stimuli have been used by our laboratory previously (21) and are based on Avenanti et al. (20). The videos were recorded using a JVC handycam recorder and edited using Adobe Premiere Elements software. The task was presented to participants using e-prime software (version 2; http://pstnet.com). Participants were instructed to pay attention to the stimuli but no further instruction were given.

**Fig. 1.**
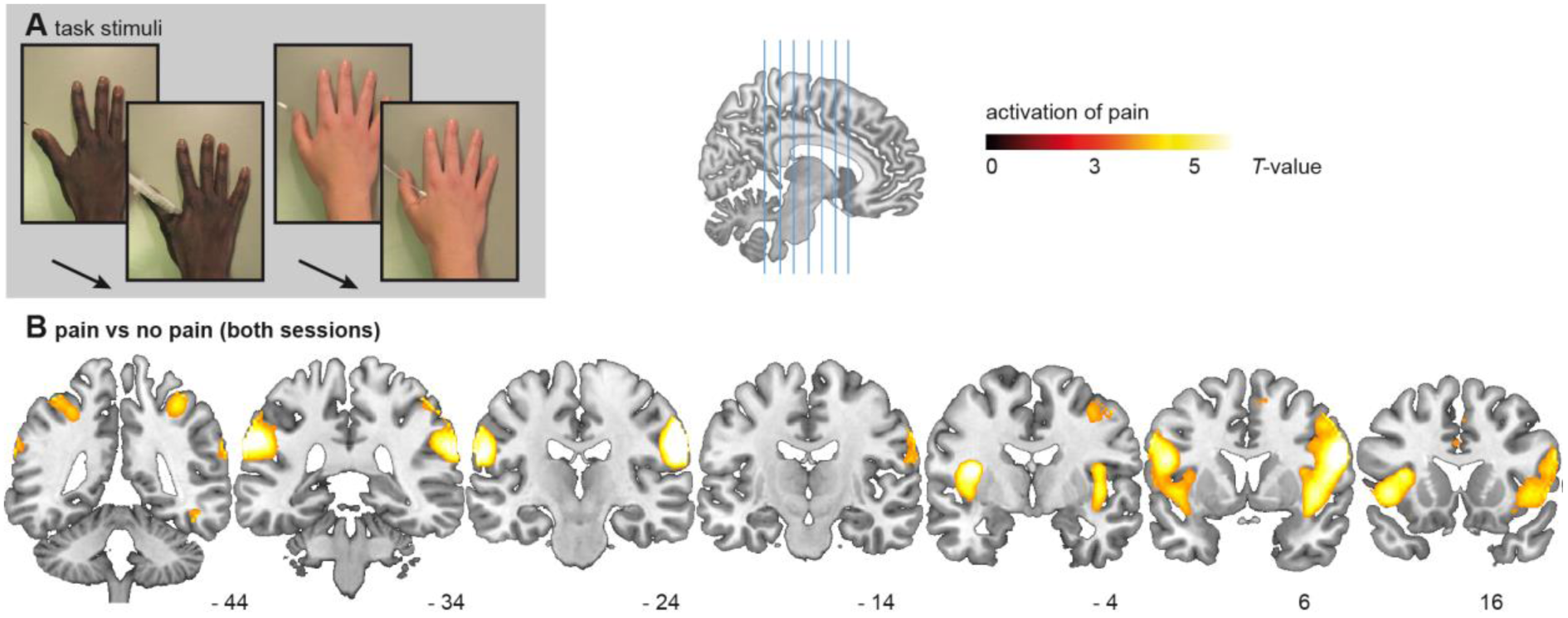
**A)** Pictures from the empathy for pain task. From left to right: the black hand in the pain condition, and the white hand in the no-pain condition. **B)** Coronal brain slices of the T-map for the contrast of pain versus no-pain in the combined placebo and testosterone conditions, overlaid onto a T1-weighted canonical image. Accompanying MNI-coordinates on the Y-axis are presented below. The T-map is thresholded at P < 0.001 (uncorrected) for illustration purposes only.

### 2.5 fMRI scanning parameters

All scans were obtained using a 3 Tesla Magnetom Allegra Siemens dedicated head MRI scanner (Siemens Medical Systems GmBH, Erlangen, Germany) with a four-channel phased array head coil, at the Cape Universities Brain Imaging Centre. Whole brain T2* weighted 2D-echo planar imaging (EPI) functional volumes were acquired with 36 ascending axial slices. The following parameters were used: EPI factor = 64, TR/TE:2s/27 ms, FA 70°, FOV (anterior-posterior, inferior-superior, left-right): 64*64*36 slices, voxel size: 3.5 × 3.5 × 4mm. Five volumes from start of the task were discarded to allow MR signal to stabilize, and 295 usable functional volumes were acquired. A T1-weighted high resolution structural scan (magentization-prepared rapid gradient echo; MPRAGE) was obtained once for each participant using the following parameters: TR/TE: 2.53/6.6 ms, flip angle 7°, FOV 256*256*128 mm, voxel size: 1 × 1 × 1.33mm, volume acquisition time: 8 minutes 33 seconds.

### 2.6 fMRI data analysis

MR scans were analysed using SPM8 (Wellcome Department of Imaging Neuroscience, London, UK). Pre-processing included; slice-time correction, motion correction of the 6 motion parameters and the sum of squared difference minimization, volume realignment to the middle volume and AC-PC realignment to improve co-registration. Functional and structural volumes were co-registered and subsequently normalized to standard (MNI152) space using an indirect normalization procedure (24) and resampled into 4mm isotropic voxels using 4th degree B-spline interpolation. Finally, all images were smoothed using an 8 mm FWHM Gaussian kernel, which addresses residual between subjects variance.

Statistical analysis of fMRI data at the individual subject level was performed within the general linear model framework. Neural responses to the presentation of the stimuli were modelled using 2.5s boxcar function convolved with the hemodynamic response function, and these were included as regressors for the four task conditions. They were white/pain; black/pain; white/no-pain; black/no-pain. Six rigid body transformation parameters obtained during realignment were also included as regressors. High pass filter cut-off was set at 1/128 Hz. For each participant, contrast maps were generated for the main effect of the separate conditions versus rest across sessions which were then entered in the second level model.

Second level random effects modelling tested the null hypothesis of zero difference across participants between the drug and placebo conditions. Whole brain and ROI analyses were run using FWE corrected voxel level significance, thresholded at p < 0.05 throughout. A three factor full factorial 2×2×2 ANOVA (drug/placebo, pain/no-pain, black/white) was modelled using event-related onset times of the task stimuli and tested for whole brain and ROI effects. In addition, to control for the possibly confounding effects of session order, 2D:4D, and skin colour of participant, these factors were entered as a covariate in three within-subjects repeated measures ANOVAs. Of the 24 participants, 10 received the drug in their first session, and 14 received the placebo in their first session; this asymmetry called for session order to be considered carefully. After all four ANOVAs were run, main effects of drug were tested, as well as main effects of pain, stimulus colour, and the interactions between the three factors. Since the chosen statistical threshold (FWE corrected at an alpha of .05) can be considered conservative with regard to often subtle effects of endocrine manipulation, we also ran exploratory analysis at a lowered threshold of a FWE corrected alpha of .1. For visualisation of results, statistical parametric maps have been superimposed onto a high resolution canonical T1 scan with thresholds set at p < 0.001.

### 2.7 ROI masks

ROI analyses were performed based on existing knowledge of neural regions associated with empathic responses to pain, as well as those regions typically experiencing dissociating effects of T administration. The predefined anatomical ROIs used were the following: Anterior insula cortex (AIC), as used in Montoa et al. (25); supramarginal gyrus, created using the automated anatomical labeling template (AAL, 26); amygdala, based on the probabilistic atlas from Amunts et al. (27) as implemented in SPM’s anatomy toolbox (28). Additionally, peak coordinates for the medial ACC (ACC; x = -2, y = 23, z = 40) and pre-supplementary motor area (pSMA; x = 6, y = 18, z = 48) were taken from a meta-analysis on pain perception (15), and 6mm circular ROIs were created in Marsbar (29). The peak voxel threshold FWE p < 0.05 was used for all ROIs. Additionally, signal beta estimates were extracted for five bilateral ROIs using Marsbar (29). The beta estimates for the AIC, amygdala, ACC, preSMA, and supramarginal gyri were entered in a repeated measures ANOVA to test for additional analysis on the main effects of drug and task conditions.

## 3 RESULTS

### 3.1 Main effects of task

Overall, the main effect of the pain stimuli compared to the control stimuli resulted in robust activation of the regions incorporated in the ‘pain matrix’. In the whole brain analysis we observed increased activation in regions associated with visual processing and detection of bodily features, including the bilateral supramarginal gyrus, bilateral middle temporal gyrus, bilateral inferior frontal gyrus as well as bilateral insula (see Table 1 and Figure 1B). ROI analyses for the same contrast detected additional activation in the medial ACC, and the preSMA. For the opposite contrast, of control stimuli versus pain, regions that activated were the bilateral middle frontal gyrus, left posterior cingulate cortex nearing the left precuneus, and right cuneus, all at the whole brain level. For the main effect of hand colour we first tested activations in response to viewing black hands versus white hands and observed activated clusters in the right fusiform gyrus, right middle occipital gyrus, right lingual gyrus at the whole brain level, as well as a small cluster in the right amygdala. A few voxels within the supramarginal gyrus showed stronger activation for the opposite contrast of white versus the black (Table 1). Critical to the aim of the current study was the contrast comparing T with the placebo condition or the interactions of drug with the other factors. However, no main effect of drug, or interaction with the other factors reached significance.

Including participants’ skin colour, 2D:4D, or session order as a covariate in the full factorial did not alter the results. Our more exploratory analyses with an alpha set at .1 (FWE corrected) neither showed effect of the factor drug administration, or interactions of drug with another factor.

**Table 1:**
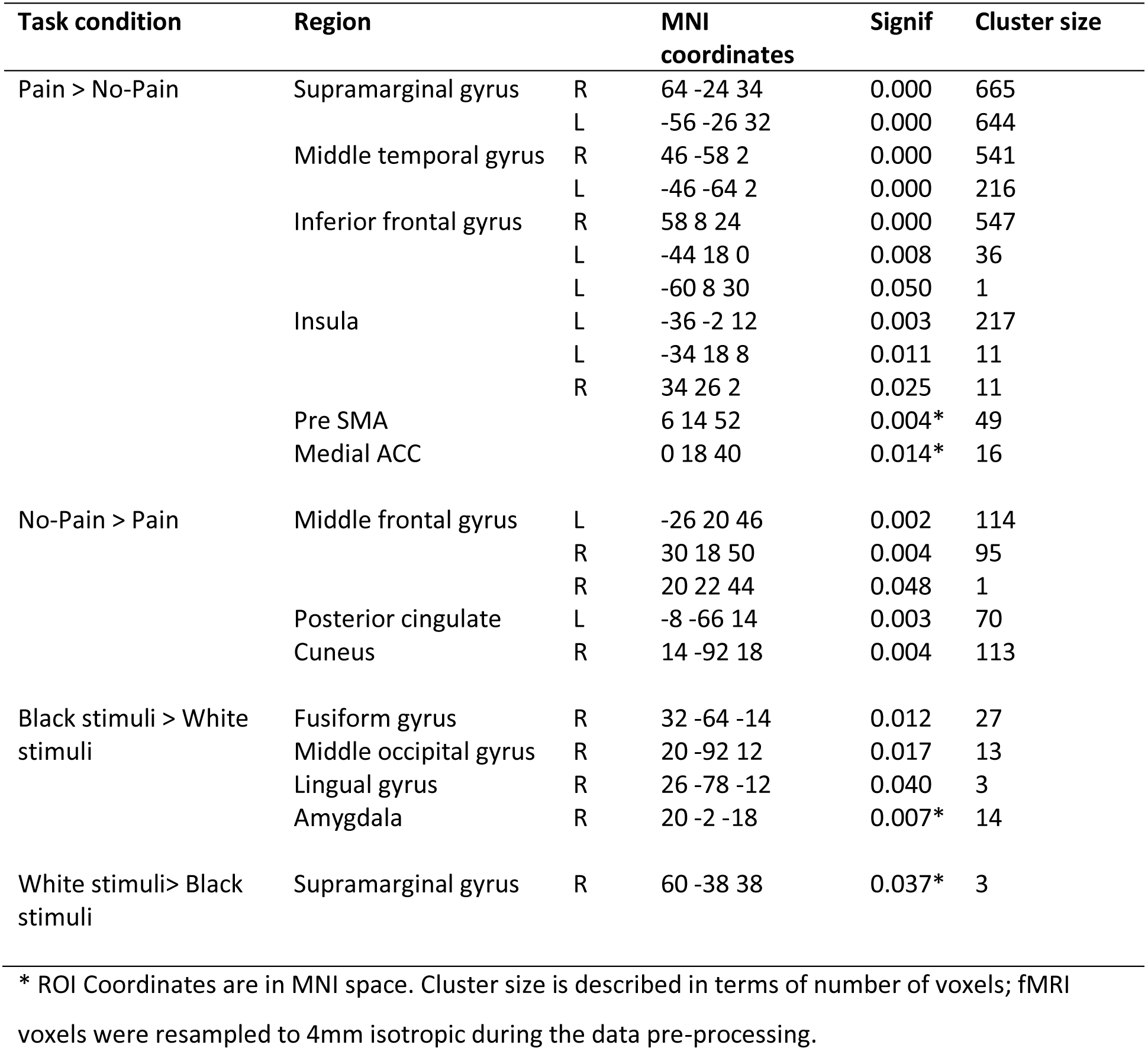
fMRI effects of task and drug.

### 3.2 Additional analyses on extraction of the ROI

The analysis of the extracted beta values of the RIOs confirmed the whole brain analyses. The AIC (p = 0.003, f = 10.704, df = 1,23), preSMA (p = 0.012, f = 7.410, df = 1,23), and supramarginal (p < 0.001, f = 22.114, df = 1,23) responded significantly in the pain condition. However, no other conditions or interactions were significant. Finally, as a control analysis, we ran an additional ANCOVA with 2D:4D ratio as a covariate to test for possible interactions of drug administration with the 2D:4D ratio, and with participant skin colour as a between subjects factor to investigate the effect on differential responses to pain depending on stimulus skin colour. However, the added factors did not significantly interact with the effects of interest.

## 4 DISCUSSION

As a main finding of the experiment, the current paradigm showed robust activation of regions associated with pain processing when witnessing others in pain such as the insula, ACC, and pre-SMA. These results are in line with previous studies using stimuli that elicit neural empathic responses to pain in others (21,17,30). Most importantly however, our study investigated the effect of T administration on the observed neural responses towards pain in others. In our current within-subject sample of 24 participants, no such effects were found.

Based on previous studies that have demonstrated downregulating effects of T on empathic behaviour (8,10,9), it was hypothesized that T would attenuate neural responses to pain in others. Also, it was predicted that similar to the effects of other endocrine factors such as oxytocin, the effect of T might differ between different aspects of empathy (21). However, no effects of T on neural responses to pain in others were observed. An important notion with regard to these findings is that the previous behavioural studies measured cognitive empathic behaviour (‘mindreading’, 9,10) and automatic facial imitation (8), but not empathy for pain in others. Thus, it is currently unknown whether T alters behavioural responses to pain in others, and the exact relation between neural responses to others’ pain in the pain circuitry and behavioural responses is also unclear. As described above, oxytocin can under some conditions increase empathic subjective responses to pain in others (31), yet this neuropeptide strongly decreases neural responses in the pain circuitry when seeing pain in others (21). It could be that the observed neural responses towards pain in others reflect aversiveness and personal distress (32,33), which if reduced by oxytocin can result in more appropriate helping behaviour (34). Relating to our current findings, it might thus be that T does not affect aversive responding or experienced personal distress. This interpretation highlights the importance of publishing null-findings, especially in the light of the current criticism on the possible publication bias in oxytocin administration studies (35). Future studies combining T administration with sensitive measures of behavioural responding to pain in others should however test the validity of our interpretation.

Given that previous studies have observed robust changes (indexed by medium-to-large effect sizes) in neural activation after the same T administration as currently used in within-subject designs with smaller samples (36,37) we can be confident that our current study setup has the power to detect effects of similar magnitude. That being said, our current sample size is less sensitive to detect small, or small-to-medium effect sizes, and thus replication of this null-finding with larger samples is needed to exclude more subtle effects of testosterone on the neural correlates of pain observed in others. Also, recent work on oxytocin has shown differences in the effects of drug administration depending on whether the study followed a within- or between subject design, and showed more robust effects of drug in the between subject design (38). Since the current study follows a within-subject design, this could also be taken into account in future studies.

Irrespective of the effect of drug, we did encounter effect of hand colour of the stimuli on our data. Relative to stimuli of white hands, black hand stimuli activated the lingual gyrus and fusiform gyrus, two regions that are well established to be related to visual processing of human forms (39,40). Fusiform gyrus activation noted in this contrast has also been noted in work that has found increased activation when viewing visuals of ingroup members (41,42). In addition, we observed increased activation of the amygdala for the black versus white hands. Although amygdala activation has been related to ingroup bias (43), this is unlikely the case in our current study. First, the effects of skin colour of the stimuli did not interact with the pain condition, and were thus similar for the pain and no-pain condition, more likely reflecting differences in visual processing. Furthermore, the effects of colour of the stimuli were unrelated to participants skin colour. Our sample was purposefully selected to consist of participants of different ethnicity to increase the generalizability of the findings (black: n = 13; white: n = 8; mixed ethnicity: n = 3), and this also allowed us to test whether participants’ skin colour predicted the effect neural responses to images of black or white hands in pain. However, extracted data from the ROIs showed no interaction between participant colour with pain that depended on the skin colour of the stimuli, indicating that the neural responses of students of different ethnicity to the black and white stimuli was similar. Thus, our current data does not replicate previous studies that found differential effects towards pain in others depending on race (18–20). Cultural differences between European and Asian students and those born into a post-apartheid South Africa might account for these differences. It must however be noted that the current study was not set up to investigate ethnicity biases in empathic responding to pain in others but focused on the effect of T, as the groups were not balanced for participant ethnicity, and the current sample size limits the power to detect subtle effects of ethnicity on the data.

In conclusion, corroborating previous findings, robust task effects were observed in brain regions involved in the processing of pain when observing pain in others. This activation pattern was however unaffected by our placebo-controlled T administration. Since neural responses towards pain in others might particularly signal aversiveness and experienced personal distress (32,33), it could be that T does not affect this particular aspect of empathic responding, in contrast to the downregulating effects T has on cognitive empathic abilities (9,10).

